# Detection of bifidobacterial lipoproteins with anti-viral potential in donor human milk

**DOI:** 10.64898/2026.07.21.739262

**Authors:** Siwar Ayari, Famara Sané, Frank Piva, Stéphanie Devassine, Fabrice Bray, Philippe Gervois, Marie-Bénédicte Romond

## Abstract

**Background:** Group B coxsackieviruses (CVBs) are involved in triggering type 1 diabetes. Free bifidobacterial lipoproteins (BLps) prevent CVBs ce **l** infection. Our objective is to isolate free BLps potentially released by bifidobacteria in breast milk and document their bioavailability.

**Methods:** *Bifidobacterium breve* and *B. longum* were quantified by qPCR in samples donated to the hospital’s biobank (DHM) within two weeks following delivery. BLps were captured onto CV B4, analyzed by SDS-PAGE and competitive ELISA. BLps transport through Caco-2 monolayers was monitored after DHM contact. Titration of anti-CV B IgA was carried out by ELISA.

**Results:** Among the 90 enrolled donors, seven were excluded, 68 donated a unique sample and 15 donated multiple specimens (2 to 12). *B. breve* and *B.longum* were detected in 93.0% and 32.8% of unique DHM samples, respectively. The two species showed unstable counts in the mult ip le donation group. Although at highly variable amounts, BLps were readily detected. Free *B.longum* Lps were found in absence of *B.longum* itself. The BLps crossed the ce **l** layer within 4h still binding CV B4.

**Conclusion:** DHM contained BLps able to cross a ce **l** layer mimicking the intestine sti **l** retaining their capacity to bind CV B. It suggested a possible newborn’s systemic protection against CV B4 infection that needs further investigation.

## INTRODUCTION

Type 1 diabetes (T1D) is a chronic autoimmune condition characterized by hyperglycemia with long-term insulin dependency [1]. Both genetic susceptibilities, mainly defined by the HLA class II gene region, and environmental factors are crucial in the development of islet autoimmunity and T1D [2]. Clinical and epidemiological studies show that infections with Coxsackievirus B (CVB) and other enteric viruses (EV) can precipitate autoimmunity in T1D-susceptible individuals. CVB, which can be found in the pancreatic islets of most patients with T1D, can accelerate disease progression by activating the immune system [3].

Enteroviruses are non-enveloped, single-stranded icosahedral RNA viruses classified within the *Picornaviridae* family that primarily display faecal–oral transmission, with occasional cases of vertical and respiratory transmission. EV spread and replicate via the upper respiratory and gastrointestinal tracts and invade the beta islet ce **l**s via the coxsackievirus adeno receptor (CAR) [4]. Anti-CVB antibodies are detected in human milk (HM) but the yield depends on the mother’s exposure to EV [5].

Recently, we showed that lipoproteins released from the ce **l** wa **l** of two bifidobacterial species prominent in childhood prevented the CVB4 ce **l** infection by mimicking CAR and blocking the virus entry [6]. Although the HM microbiome is known to harbor bifidobacteria, there is sti **l** no evidence proving that the HM holds ce **l** free bifidobacterial Lps (BLps) [7]. BLps structure with fatty acid moieties linked to the protein sequence hampers an easy detection in complex matrices [8, 9].

To overcome the chemical complexities, we developed an affinity column using CVB4 to capture BLps. Investigation of BLps occurrence in HM samples donated at the biobank was designed in two steps: (i) quantification of HM colonization with two BLps secreting bifidobacterial species, *Bifidobacterium longum* and *B. breve*, in HM donated (DHM) samples, (**i**) detection of BLps in multiple DHM harboring or not bifidobacteria. The present study also explored the possible delivery of BLps to internal organs following DHM intake.

## Material and Methods

### Study Design and Participants

We applied a prospective cohort design. The samples used in this study were collected from 2013 to 2017 and subsequently stored until analysis [5]. Lactating women were recruited by the medical staff at the biobank of Lille Hospital among mothers wi **l**ing to donate their milk. At enrollment, participants completed a study questionnaire to collect demographic information and a history of antibiotherapy. According to the biobank rules, participants were instructed to self-collect human milk into a clean, sterile container at home using an electric or manual pump. The milk was frozen in the participant’s home freezer then transported to the biobank where a fraction (at least 20 mL) was sampled, transferred to the laboratory and kept frozen at -20 °C until treatment. The study was approved by the regional committee for medical research ethics and written informed consent was obtained from each participant.

### Enumeration of Bifidobacteria

As shown in Fig. 1, milk samples (20 mL) were thawed, centrifuged at 19 000 g at 4 °C for 30 min. The lipid layer was removed and both supernatant and pe **l**et were collected. Total DNA from the pe **l**et was extracted using the Nucleospin Tissue kit (Macherey Nagel, Hoerdt, France). *Bifidobacterium longum* and *B.breve* DNA were quantified by qPCR [10]. Briefly, qPCRs were set out in 20 μL final volumes containing 5 μL of DNA template (10 to 100 ng), 4 μL of Evagreen x5 Mix (Euromedex, Souffelweyersheim, France) and 2 µL of each primer (*B.longum* FW: CCA GCT GCT GCT TGA GTT C, RV: CCA GCT GCT GCT TGA GTT C; *B.breve* FWD: GTG CTG ATC GCT TCT CAG G, RV: CAT CTC CTT ACG CCG GTT GT). The amplification conditions were 12 min denaturation at 95°C, followed by 40 cycles of 15 s denaturation at 95 °C, 20 s for annealing at 63 °C for *B.longum* and 60 °C for *B.breve* and elongation for 30 s at 72 °C (Mastercycler realplex, Eppendorf). PCR products from samples and isolated bacteria were controlled by melting curve analysis and migration on 2% agarose gel (Eurogentec, Liège, Belgium). The qPCR assays were calibrated using known amounts (from 0.01 fg up to 10 pg, i.e., from 42 to 4.2×10^7^ copies) of PCR-amplified target gene fragments from corresponding bifidobacterial species. [11] *Bifidobacterial Lipoprotein (BLps) isolation and analysis*

**Figure 1:**
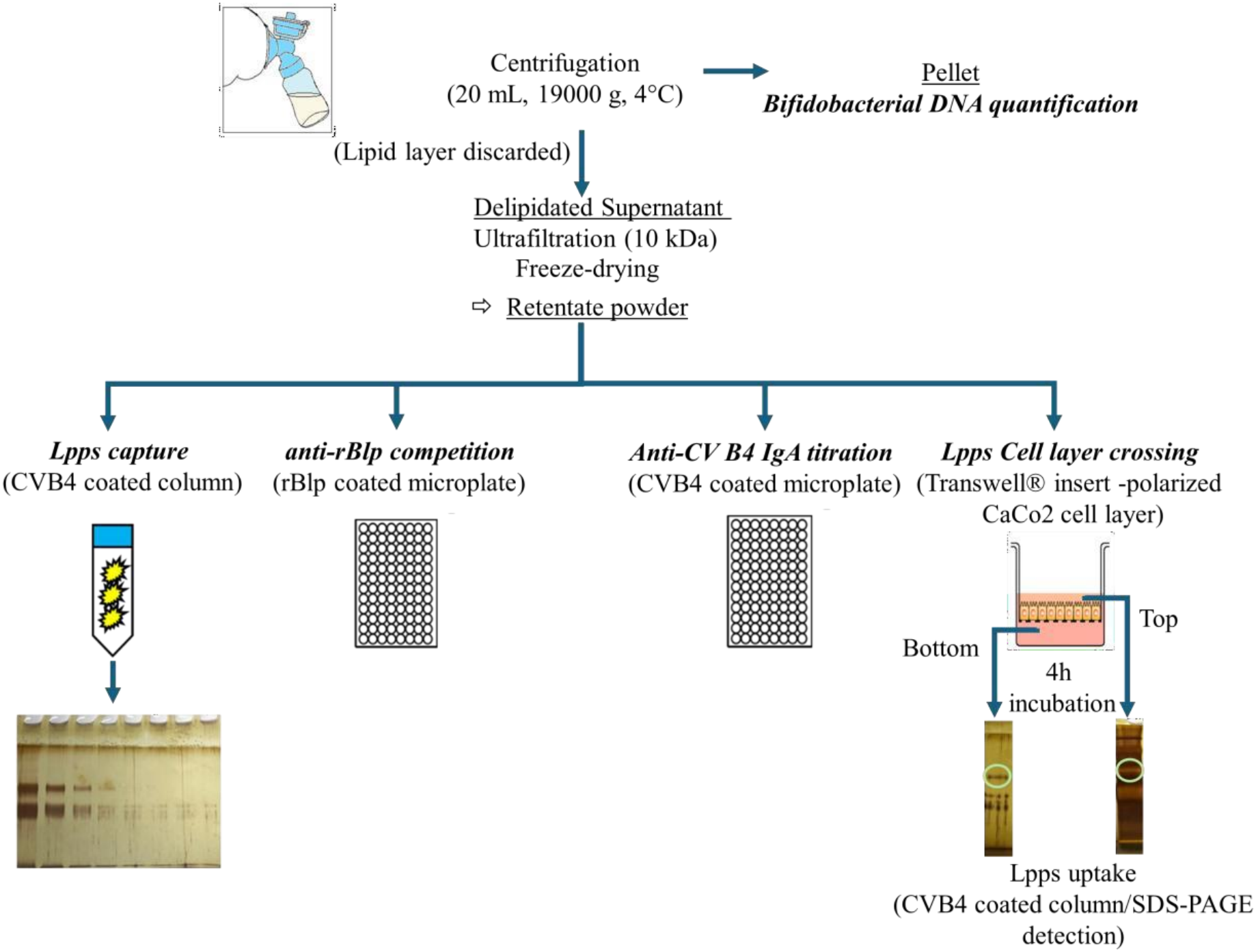
Isolation and analysis of bifidobacterial lipoproteins from delipidated breastmilk powder devoid of oligosaccharides and enumeration of *Bifidobacterium breve* and *B. longum*.

Breastmilk supernatant prepared as mentioned above was ultrafiltrated using a 10 kDa cut-off threshold membrane (Vivaspin, Sartorius, France) to remove lactose and polylactosamines. The retentate was further diafiltrated and then lyophilised. The binding of free bifidobacter ia l lipoproteins (BLps) to Coxsackievirus B4 covalently immobilized onto Aminolink resin (Thermo Fisher Scientific, Vi **l**ebon-sur-Yvette, France) was carried out by injecting 800 µL of lyophilised Human milk powder (HMP) reconstituted in 0.1 M PBS buffer, pH 7.0, at a concentration of 5 mg/mL into the column. BLps were eluted using 0.1 M glycine, HCl, pH 2.8. The collected sample was neutralized immediately, extensively diafiltrated against sterile water into a 5 kDa cut-off membrane (Vivaspin) and concentrated up to 50 µL. BLps were separated by 10% SDS-PAGE and stained by silver staining [9].

The quantification of BLps was carried out using serial dilutions of the eluted fractions. Known amounts of BLps isolated from fermentation with *B.longum CBi0703* (from 1 ng up to 1 µg) were used to determine the lowest amount detected by silver-stained SDS-PAGE (i.e. 50 ng per lane). Bands around 80 kDa were further probed by Orbitrap MS following ce **l**ulase and trypsin digestion as we **l**. Peptides were identified by Proteome Discover software against *B.breve* C50 and *B.longum* CBi0703 Lps sequences. Search for O-and N-glycan derivatization was carried out using Byonic software.

The lowest dilution of the eluted fraction with a specific BLps band was considered as containing 50 ng/volume laid onto the gel. The final equation was:

Amount BLps (ng/mg HMP) = [(50 ng/ gel vol^a^) x dilution x total eluted vol)]/dry milk extract weight laid onto the CV B4 affinity column

(^a^: volume laid onto the gel= 8 µl)

Furthermore, HMPs were subjected to an in-house competitive ELISA assay. Nunc 96-well plates were coated with 1.25 ng recombinant protein of *B. longum* Lps (rBlp) diluted in coating carbonate solution (40mM, pH 9.6) overnight at 4 °C. The polyclonal anti-rBlp IgG diluted in 8mM PBS, 1 % BSA, 0.05 % Tween-20 (1:1250) was incubated for 1 h at 37°C (V/V) with HM retentates ranging from 12.5 to 100 µg HMP /ml (i.e. 6.25 to 50 µg HMP and anti-rBlp IgG at 1:2500 in one ml). The recombinant protein at concentrations ranging from 12.5 to 100 ng rBlp/ml were incubated para **l**elly at 37°C for 1 hour with anti-rBlp IgG diluted in 8mM PBS, 1 % BSA, 0.05 % Tween-20 (1:1250) (i.e. 6.25 to 50 ng rBlp and anti-rBlp IgG 1:2500 per ml).

The incubated mixes (200 µl) were then added to the microplates coated with 1.25 ng rBLps. After 1h incubation at 37°C, the microplates were washed three times before adding biotinylated anti-rabbit IgG diluted at 1:1000. Streptavidin diluted at 1:1000 was incubated for 1h at 37°C after 3 washes. SigmaFast™ OPD reconstituted in water was added to the we **l**s for color development, and the reaction was stopped by adding a 2 % sulfuric acid solution. Plates were read at 490 nm. A standard curve with increased coated rBlp (from 0.0195 up to 20ng) with direct incubation with anti-rBlp IgG diluted in 8mM PBS, 1 % BSA, 0.05 % Tween-20 (1:2500) was used to determine the linear range of OD values and to monitor the reproducibility between the in-house ELISA assays.

The linear range of OD values was comprised between 1.56 to 2.5 ng coated rBlp. The observed inhibition of anti-rBlp 1:2500 with 6.25 ng/ml was 52 ± 4 %, which corresponded to the expected 50% inhibition. But the competition with 12.5 ng/ml was less efficient with only 41 ±2 % inhibition, which contrasted with the expected 100%. The highest concentrations of rBlp (25 and 50 ng/ml) did not achieve >50% inhibition rate (36 ± 9 and 24 ± 20%, respectively). The poorer inhibitory effect at high concentration reflected the hydrophobic nature of rBlp resulting in the aggregation of the monomeric forms when increasing the concentration in hydrophilic solvents [9]. Therefore, inhibition of less than 45% is to be disregarded. A positive detection of *B.longum* protein sequence in HDM samples wi **l** thus be associated with a threshold of at least 46% inhibition of anti-rBlp binding to the coated rBlp.

### Anti-CV B4 IgA quantification

An in-house assay was used to measure CV B4 specific breastmilk IgA. Nunc 96-we **l** plates were coated with 10^4^ pfu CV B4 (E2 strain) diluted in coating carbonate solution (40 mM, pH 9.6) overnight at 4 °C. Plates were washed three times (8 mM PBS, 0.05 % Tween-20, pH 7.4) and incubated for 1h at 37°C with blocking solution (8 mM PBS, 1 % BSA, 0.05 % Tween-20).

Anti-IgA antibodies, streptavidin and breast milk samples were diluted in PBS solution (8 mM) supplemented with Tween-20 (0.0375%) and BSA (2%). HMP were added to we **l**s in duplicate and incubated overnight at 37 °C. Plates were then washed three times and incubated for 1h at 37 °C with a biotinylated anti-IgA antibody at a 1:2500 dilution, prior streptavidin/OPD color development. Plates were read at 490 nm. OD values within the linear range of a standard curve (breast milk pool exhibiting the highest neutralizing potency) were used to interpolate the concentration of CV B4 binding IgA antibodies.

### Transport of BLps across the Caco-2 cell layer

To study BLps capacity to pass across the intestinal epithelium, confluent polarized Caco-2 cells were used as a barrier for particles crossing from the Apical to Basolateral chambers.

Caco-2 ce **l**s were cultured in Dulbecco’s Modified Eagle’s Medium (DMEM), supplemented with 10% fetal bovine serum, 1% nonessential amino acids, and 4 mM glutamine without antibiotics in a humidified incubator at 37°C and 5% CO^2^. The cultures were seeded onTranswells ® (0.4 μm pore size, 1 cm^2^ growth area; Corning Costar Co. USA) at a ce **l** density of 3× 10^5^ ce **l**s/membra ne. The ce **l** monolayer was fed with fresh medium every 2 days and ce **l**s were used on day 21 for the transport experiments [12]. DMEM was replaced by Earle’s balanced salt solution (EBSS) (Gibco, ThermoFischer Scientific) as the transport medium. The ce **l** layer was carefully rinsed with pre-warmed HBSS. To prevent any bacterial contamination and reduce ce **l** stress to the minimum, the membrane integrity was monitored by incubating 400 µL DMEM (comprising the neutral red dye that can cross the Transwell insert if the ce **l** layer is deteriorated) in the apical chamber for 15 min at 37 °C and measuring deviation in the HBSS buffer from the basolateral chamber to the optical density of a control HBSS buffer at a wavelength of 700 nm (red wavelength). After rinsing the ce **l** layer, HMP reconstituted in HBSS at various concentrations was added to the upper chamber and incubated for 4 h. Known amounts of BLps isolated from fermentation with *B.longum* CBi0703 were used to determine the minimum concentration and the incubation time required to detect a transfer in the basolateral chamber. Samples were withdrawn from the apical and basolateral compartments. After being concentrated up to 50 µL, they were diluted in 0.1 M PBS buffer (400 µL) and injected into the CV B4 affinity column. Elution and identification were performed as described above.

## Results

### Collection of HM samples

Ninety mothers were enrolled. Samples from 7 mothers were excluded due to uncertainty about the date of birth (4), donation after the recruitment period (1), and failing to express suffic ient volume (2). Sixty-eight mothers donated a unique sample, and 15 ones donated between 2 and 12 samples. A total of 142 samples were collected.

### Bifidobacteria in HM donations

Most of the unique donations were collected during the first week following childbirth (59 out of 68 donors). Donations were rare during the very first days after birth, with only two donations in the hours following delivery and three on day 1 (D1) after parturition. During this period, *B.longum* DNA was detected in a single sample in the hours following delivery, whereas *B.breve* DNA was commonly found (Fig. 2). On the second and third day after childbirth, the high prevalence of *B. breve* is sti **l** observed in the donations (92 and 89%, respectively), which contrasts with the low prevalence of *B. longum* (33 and 21%). The prevalence of *B. breve* and *B. longum* becomes more similar on the 4^th^ day (77 vs 69%). As i **l**ustrated in Fig.2, *B.breve* is also quantified at higher counts than *B.longum* during the first week postpartum. In the second week postpartum, only 9 unique donations were collected (2 on D10) (Table S1). The most striking observation was the lack of *B.longum* detection, whereas a **l** except one sample harboured *B.breve* DNA. In summary, 93% of unique donations harbored *B.breve* DNA, 201 whereas only 32% were positive for *B.longum* DNA within the first two weeks.

**Figure 2:**
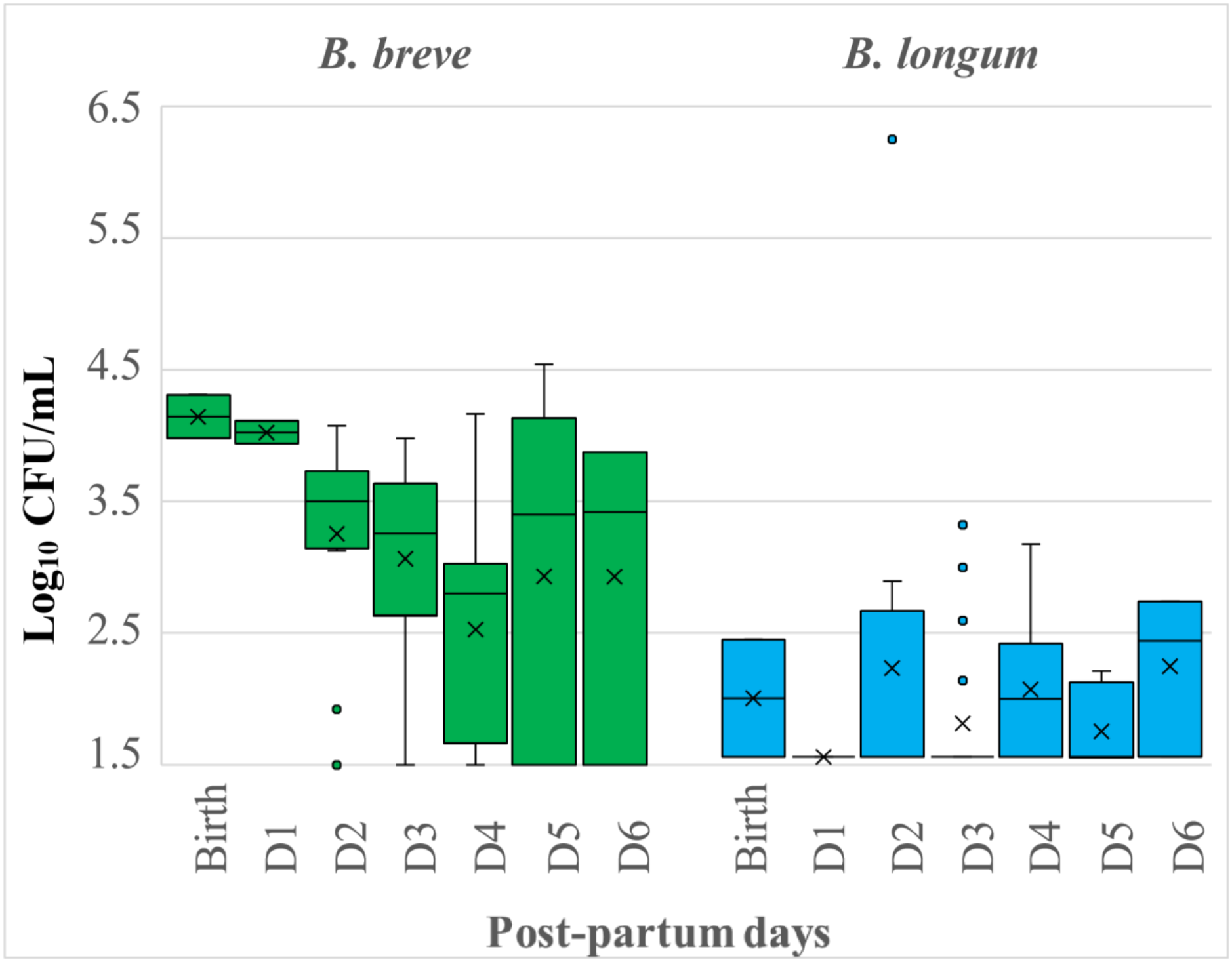
Enumeration of *B.breve* and *B.longum* by qPCR in unique donations during the first week post-delivery.

When the mothers donated several samples, *B.breve* predominance was less significant (Fig. 3). Moreover, bifidobacterial DNA from both species was intermittently detected (Fig. 3). Although the difference in prevalence was less obvious, *B.breve* was sti **l** detected in 72% of samples, whereas *B.longum* DNA was recovered in 52%.

**Figure 3:**
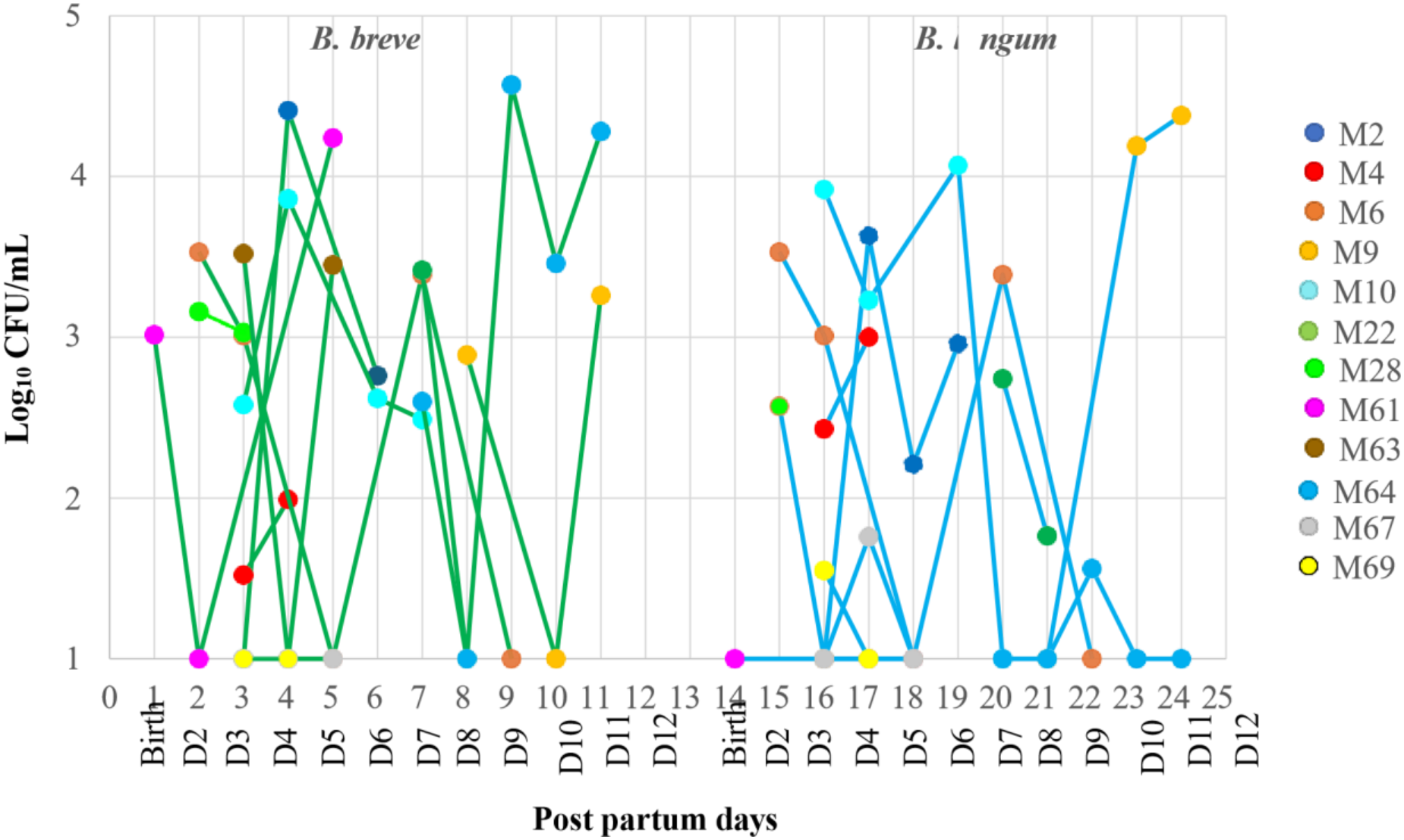
Enumeration of *B.breve* and *B.longum* by qPCR in multiple donations during the first two weeks post-delivery

### Free bifidobacterial lipoproteins detection in multiple donations

Donations from four mothers (M3, M11, M17 and M58) were further analyzed for their content in free lipoproteins. The donation period varied according to the mother. The biobank received milk donations between D2 and D5, D5 and D8, D3 and D12 and D4 and D15 from M3, M11, M17 and M58, respectively. *B.breve* and *B.longum* were detected in M17 and M58 at a significa nt level, albeit discontinuously (Fig. 4A). M3 and M11 samples were characterized by the lack of *B.breve* DNA (except for one M3 sample at D4) and a low but almost steady content in *B.longum* DNA (Fig. 4B). In contrast with the uneven display of bifidobacterial DNA, BLps were captured onto CV B4 affinity column in a **l** milk samples even when no DNA from *B.breve* and/or *B.longum* was found. As shown in Figure 4, neither the occurrence nor the amount of BLps depended on the bifidobacterial counts. Additional samples from the previous unique and multiple donations were included to further investigate the occurrence of free BLps and the secreting bifidobacteria (Table S2). The results confirmed the lack of correlation between BLps and bifidobacterial amounts.

**Figure 4:**
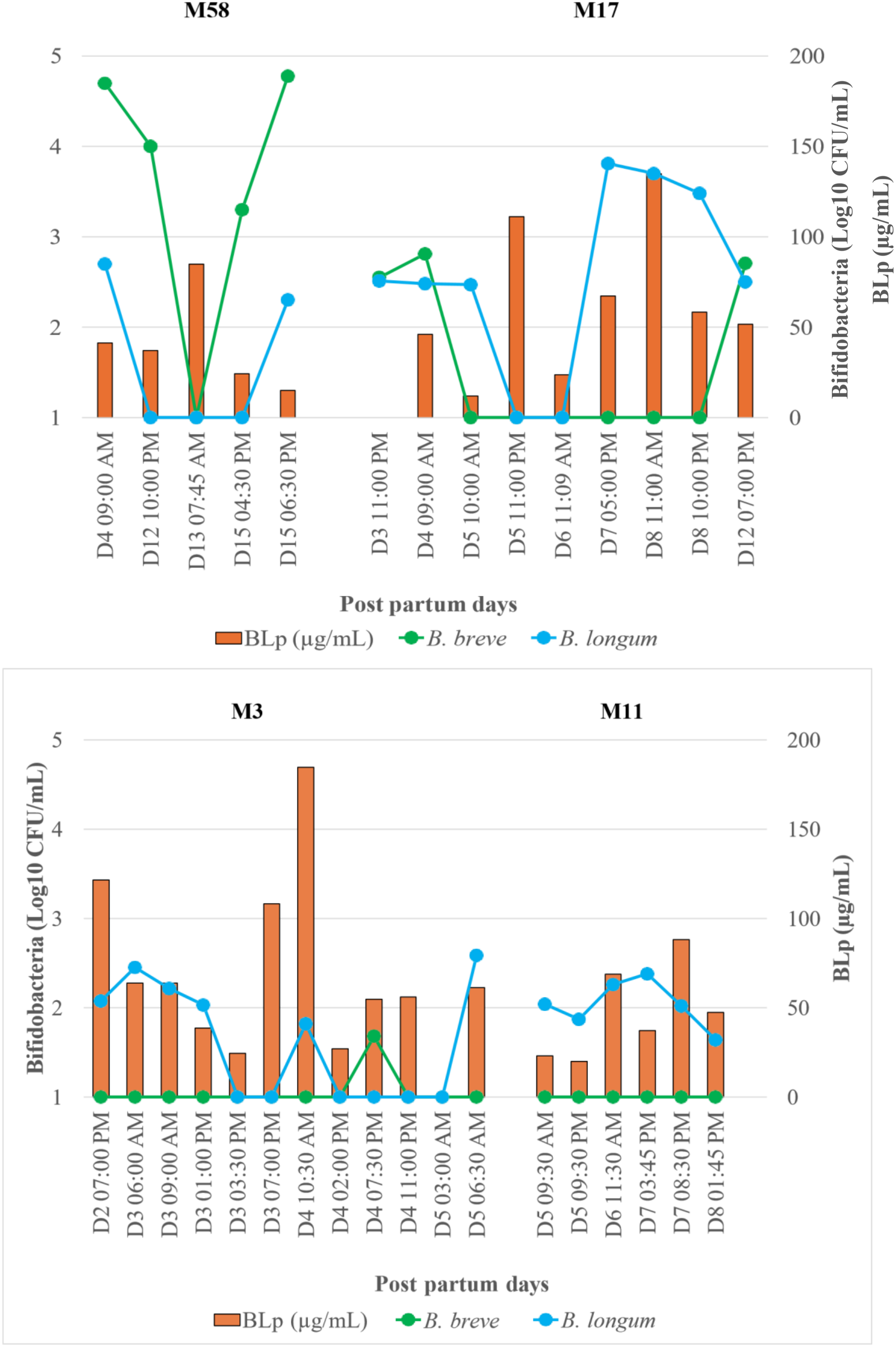
*B.longum,B.breve and* BLps in donations from M3,M11,M17 and M58.

**Figure 5:**
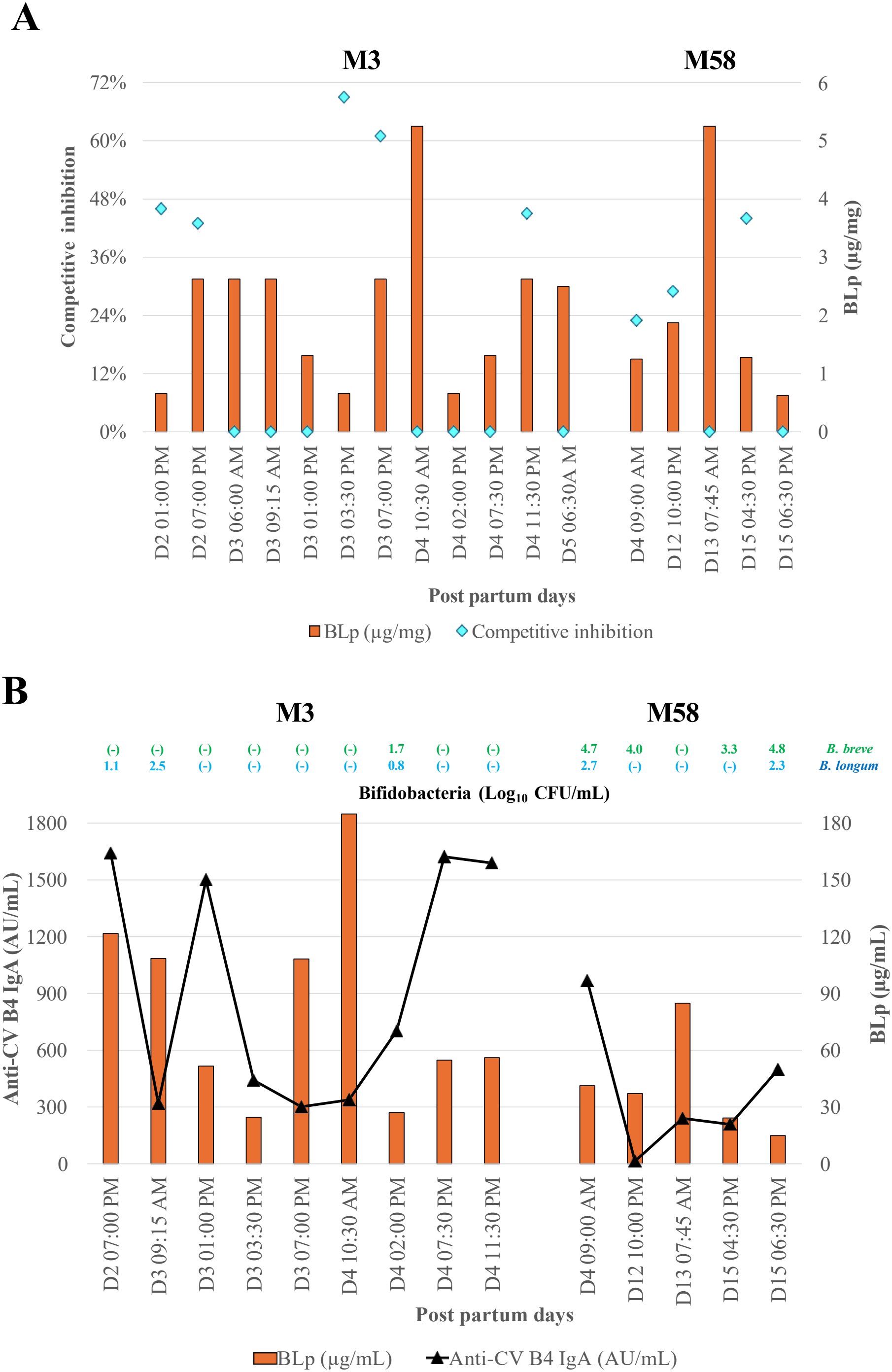
*B.longum* Lps recovery in BLps and anti-CV B4 IgA titers in multiple donations from mother 3 and 58. A) competitive inhibition using recombinant *B.longum* protein coated microplates, expressed as % of anti-rBl antibodies inhibition. B) quantification of anti-CV B4 IgA using CV B4 coated microplates, expressed as arbitrary unit/ml breastmilk.

To address more specifically the question, we determined the capacity of anti-rBlp IgG tobind to the human milk powder deprived in oligosaccharides and lipids (HMP) instead of the coated recombinant protein of *B.longum* (Table 1). Although the in-house ELISA was not sensitive due to the hydrophobic nature of the *B. longum* protein sequence (an inhibitory potency above 46% threshold being related to a positive detection of *B.longum* Lps in HDM samples), it uncovered at least 3 samples from M3 donations with a suitable inhibition rate. Among them, two samples collected on D3 at 3:30 and 7 pm did not contain *B.longum* DNA. The observations further supported that BLps detection in HM samples did not require the presence of the secreting bifidobacteria.

**Table 1:**
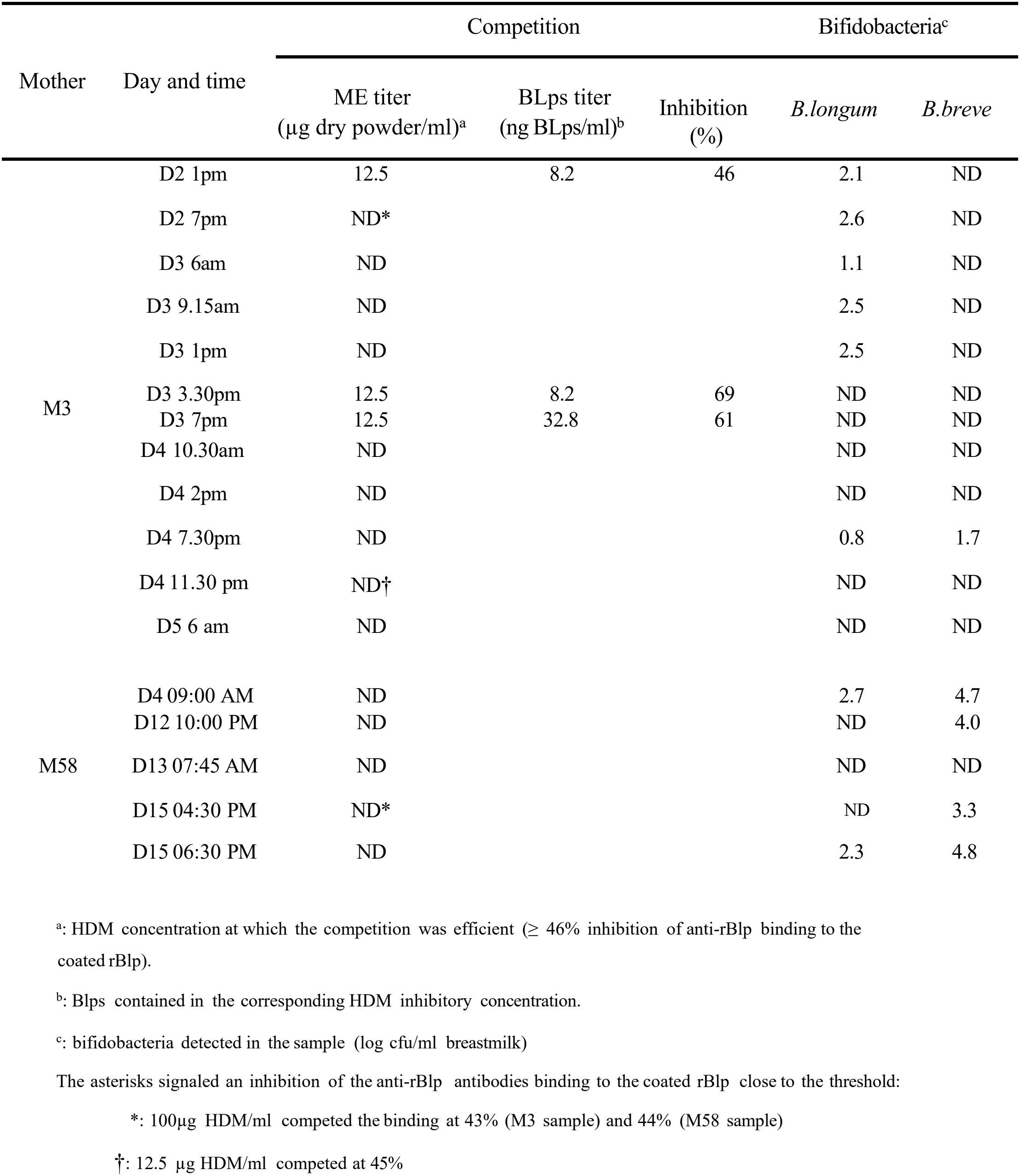
Detection of *B.longum* lipoproteins in M3 and M58 donations by competition ELISA.

| Mother | Day and time | Competition |  |  | Bifidobacteria<sup>c</sup> |  |
|---|---|---|---|---|---|---|
|  |  | ME titer<br>(µg dry powder/ml)<sup>a</sup> | BLps titer<br>(ng BLps/ml)<sup>b</sup> | Inhibition<br>(%) | *B.longum* | *B.breve* |
| M3 | D2 1pm | 12.5 | 8.2 | 46 | 2.1 | ND |
|  | D2 7pm | ND* |  |  | 2.6 | ND |
|  | D3 6am | ND |  |  | 1.1 | ND |
|  | D3 9.15am | ND |  |  | 2.5 | ND |
|  | D3 1pm | ND |  |  | 2.5 | ND |
|  | D3 3.30pm | 12.5 | 8.2 | 69 | ND | ND |
|  | D3 7pm | 12.5 | 32.8 | 61 | ND | ND |
|  | D4 10.30am | ND |  |  | ND | ND |
|  | D4 2pm | ND |  |  | ND | ND |
|  | D4 7.30pm | ND |  |  | 0.8 | 1.7 |
|  | D4 11.30 pm | ND† |  |  | ND | ND |
|  | D5 6 am | ND |  |  | ND | ND |
| M58 | D4 09:00 AM | ND |  |  | 2.7 | 4.7 |
|  | D12 10:00 PM | ND |  |  | ND | 4.0 |
|  | D13 07:45 AM | ND |  |  | ND | ND |
|  | D15 04:30 PM | ND* |  |  | ND | 3.3 |
|  | D15 06:30 PM | ND |  |  | 2.3 | 4.8 |

<sup>a</sup>: HDM concentration at which the competition was efficient (≥ 46% inhibition of anti-rBlp binding to the coated rBlp).
<sup>b</sup>: Blps contained in the corresponding HDM inhibitory concentration.
<sup>c</sup>: bifidobacteria detected in the sample (log cfu/ml breastmilk)
The asterisks signaled an inhibition of the anti-rBlp antibodies binding to the coated rBlp close to the threshold:
\*: 100µg HDM/ml competed the binding at 43% (M3 sample) and 44% (M58 sample)
†: 12.5 µg HDM/ml competed at 45%

On the contrary, HMPs exhibiting both *B.longum* DNA and BLps did not block the anti-rBlLp antibodies (M58 on D4 6:00 am and D15 6:30 am, M3 on D3 6:00, 9:30 am and 1:00 pm, and on D4 7:30 pm), which suggested that *B.longum* did not release their ce **l** wa **l** lipoproteins *in situ*. Besides, a few samples (M58 D15 6:30 am or M3 D4 7.30pm) did not display any competitive potency (below 0% inhibition). It indicated that the protein sequences of the free BLps captured onto CV B4 were specific of bifidobacterial species other than *B.longum*.

The anti-CV B4 IgA was also quantified in M3 and M58 donations using microplates coated with CV B4 (Fig.5, Table S2). No significant correlation between BLps and anti-CVB IgA was observed. However, BLps complemented the anti-CV B activity of breastmilk in several samples partia **l**y or tota **l**y deprived in anti-CV B4 IgA (Fig.5, Table S2). In these samples, BLps titres varied from 0.6 to 2.5 µg/mg HMP, corresponding to 4 to 37 µgBLps/ml breastmilk in samples deprived in anti-CV B IgA. *B.breve* and *B.longum* BLps isolated from fermentation neutralized ce **l** infection CV B4 at dosing as low as 1.6 ng/ml which further supported that the supply in HM BLps could compensate for the lack of anti-CV B IgA [6].

### Passage across the Caco-2 cell layer

BLps crossing of the ce **l** layer was studied using increased concentrations of HMP from samples donated by the mothers M3, M17 and M58 (Table 2). Prior to the assays, the amount of BLps required for detecting a passage across the ce **l** layer was determined using *Bifidobacterium longum* CBi0703 lipoproteins isolated in aggregated and monomeric forms after fermentation [9]. Increased concentrations (10, 20, 40 and 50 µg diluted in 500 µL MEM) were applied into the top compartment for 10 min up to 4 h. *B. longum* Lps were captured in the lower compartment using the CV B4 column. SDS-PAGE analysis detected *B. longum* Lps when the ce **l** layer was exposed to a minimum concentration of 40 µg *B.longum* Lps for 4 h which corresponded to a minimum of 66.7 µg BLps/cm^2^ ce **l** layer. The peptide analysis of *B.longum* CBi0703 Lps with Byonic software substantiated the O-glycan derivatization onto serine and threonine moieties, in the CHAP domain (S377, T440) as we **l** as in the helicoidal structure (S111, T119) (Fig. 6). The complex glycans comprised hexosamines and neuraminic acids such as HexNac (2)Hex(2)Neu (1 or 2 or 3). The amount of MP to incorporate into the top of the ce **l** layer was calculated according to their BLps concentration. As shown in Table 2, a few samples were applied at a lower concentration than the threshold. Two samples from M3 (i.e. on D4 and D5) were tested at a concentration of 22.7 and 50.6 µg BLps/cm^2^ ce **l** layer, respectively. No BLps were detected in the lower compartment after a 4 h contact. Increased concentrations above the threshold demonstrated BLps crossing of the ce **l** layer. Similar results were seen with samples from M58 on D5 at 6:30 pm. Crossing was detected only when the minimum BLps concentration was 66.7 µg BLps/cm^2^. An exception was seen with the sample collected on D15 at 4:30 pm. The assay was duplicated and the BLps contents were quantified in the apical compartment at the end of the 4h contact. The titers ranged between 100 and 133 µg BLps/cm^2^. It suggested that BLps were primarily in aggregated forms in MP, the monomeric forms being released during incubation which likely facilitated the passage across the ce **l** layer.

**Table 2:**
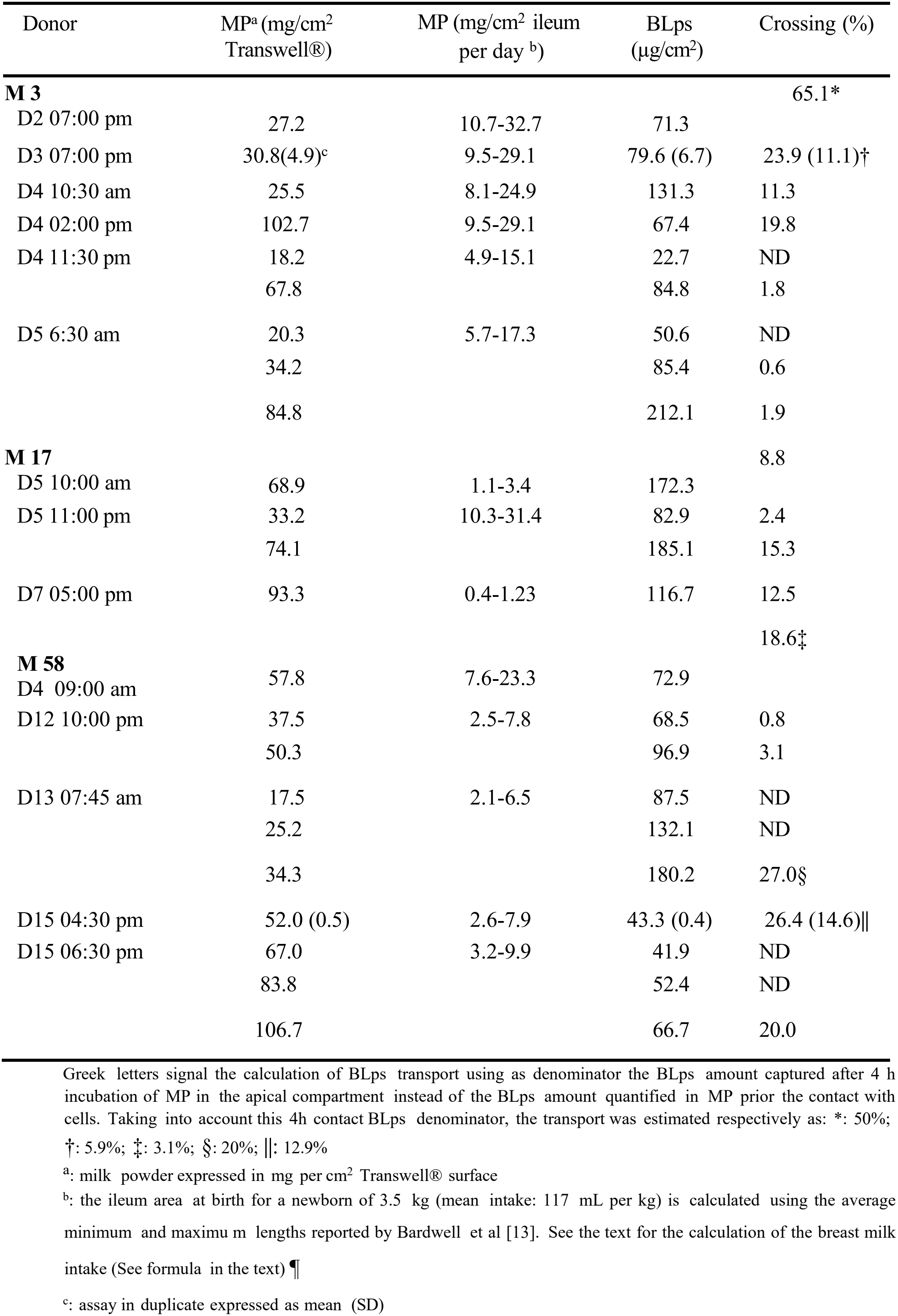
Passage of BLps across Caco-2 ce l layer.

| Donor | MP<sup>a</sup> (mg/cm<sup>2</sup> Transwell®) | MP (mg/cm<sup>2</sup> ileum per day<sup>b</sup>) | BLps (µg/cm<sup>2</sup>) | Crossing (%) |
|---|---|---|---|---|
| **M 3** |  |  |  | 65.1* |
| D2 07:00 pm | 27.2 | 10.7-32.7 | 71.3 |  |
| D3 07:00 pm | 30.8(4.9)<sup>c</sup> | 9.5-29.1 | 79.6 (6.7) | 23.9 (11.1)† |
| D4 10:30 am | 25.5 | 8.1-24.9 | 131.3 | 11.3 |
| D4 02:00 pm | 102.7 | 9.5-29.1 | 67.4 | 19.8 |
| D4 11:30 pm | 18.2 | 4.9-15.1 | 22.7 | ND |
|  | 67.8 |  | 84.8 | 1.8 |
| D5 6:30 am | 20.3 | 5.7-17.3 | 50.6 | ND |
|  | 34.2 |  | 85.4 | 0.6 |
|  | 84.8 |  | 212.1 | 1.9 |
| **M 17** |  |  |  | 8.8 |
| D5 10:00 am | 68.9 | 1.1-3.4 | 172.3 |  |
| D5 11:00 pm | 33.2 | 10.3-31.4 | 82.9 | 2.4 |
|  | 74.1 |  | 185.1 | 15.3 |
| D7 05:00 pm | 93.3 | 0.4-1.23 | 116.7 | 12.5 |
| **M 58** |  |  |  | 18.6‡ |
| D4 09:00 am | 57.8 | 7.6-23.3 | 72.9 |  |
| D12 10:00 pm | 37.5 | 2.5-7.8 | 68.5 | 0.8 |
|  | 50.3 |  | 96.9 | 3.1 |
| D13 07:45 am | 17.5 | 2.1-6.5 | 87.5 | ND |
|  | 25.2 |  | 132.1 | ND |
|  | 34.3 |  | 180.2 | 27.0§ |
| D15 04:30 pm | 52.0 (0.5) | 2.6-7.9 | 43.3 (0.4) | 26.4 (14.6)‖ |
| D15 06:30 pm | 67.0 | 3.2-9.9 | 41.9 | ND |
|  | 83.8 |  | 52.4 | ND |
|  | 106.7 |  | 66.7 | 20.0 |

Greek letters signal the calculation of BLps transport using as denominator the BLps amount captured after 4 h incubation of MP in the apical compartment instead of the BLps amount quantified in MP prior the contact with cells. Taking into account this 4h contact BLps denominator, the transport was estimated respectively as: \*: 50%; †: 5.9%; ‡: 3.1%; §: 20%; ‖: 12.9%
<sup>a</sup>: milk powder expressed in mg per cm<sup>2</sup> Transwell® surface
<sup>b</sup>: the ileum area at birth for a newborn of 3.5 kg (mean intake: 117 mL per kg) is calculated using the average minimum and maximu m lengths reported by Bardwell et al [13]. See the text for the calculation of the breast milk intake (See formula in the text) ¶
<sup>c</sup>: assay in duplicate expressed as mean (SD)

**Figure 6:**
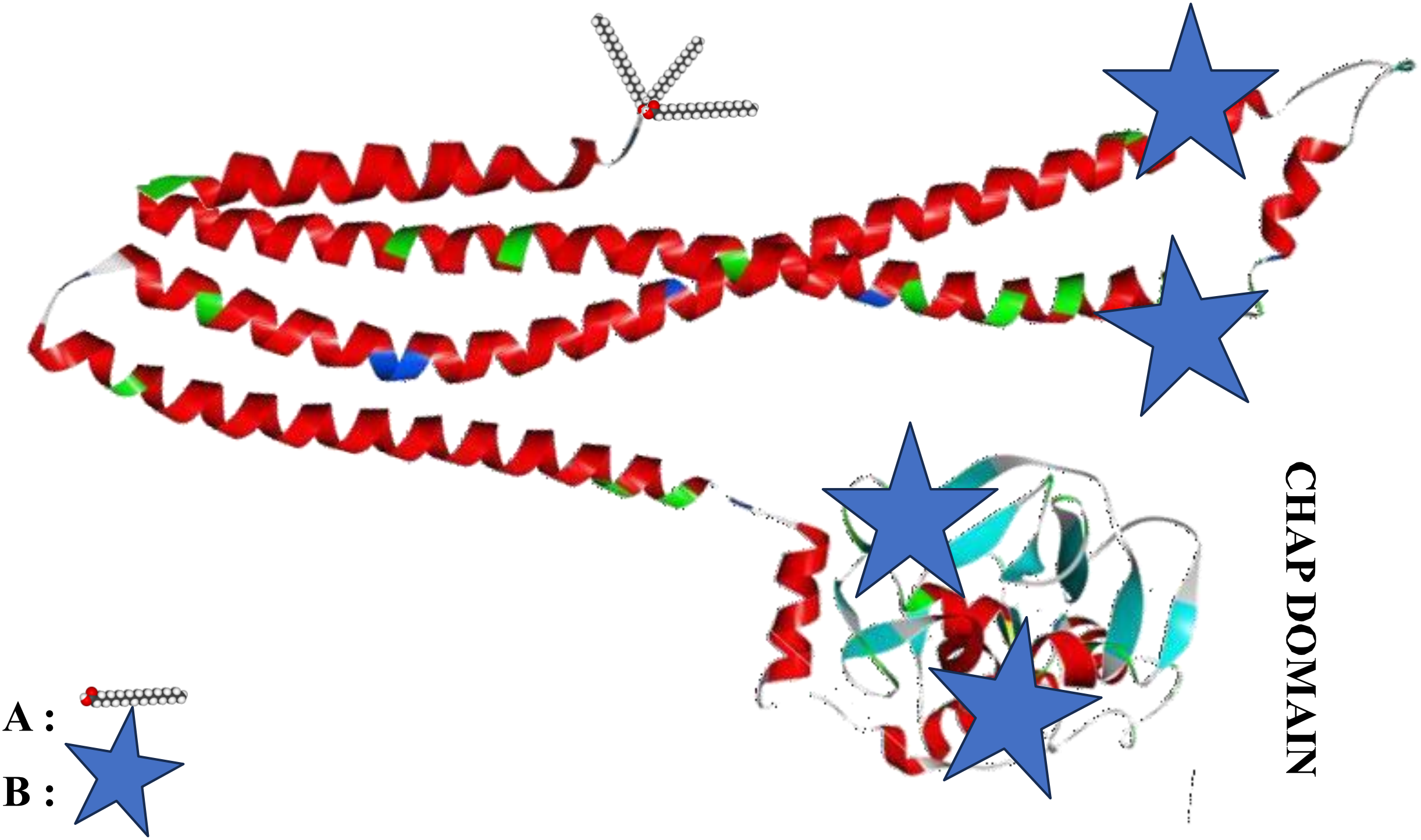
Putative conformation of BLps from *Bifidobacterium longum* with bi/triacylation (A) onto cystein in the Nterminal side and O-glycosylation on Ser/Thr (blue star (B)) detected by Byonic software on various locations of the protein sequence, including the CHAP domain.

In contrast, BLps from M58 sample collected on D13 at 7:45 am did not easily cross the cell layer. More than twice the threshold concentration was required to detect BLps in the basal compartment.

Moreover, the migration rate was not related to the BLps concentration nor the corresponding MP amount deposited onto the top compartment. For instance, the M3 samples showed the best efficiency on D2, even after correcting the estimated crossing by using the BLps titre after 4 h contact of HMP with the epithelial ce **l**s. In comparison, the sample collected on D5 at 6:30 am prevented passage even when laying almost three times the initial BLps amount.

*In vivo*, the quantity of MP in contact with the ce **l** layer is, however, constrained by the volume consumed by the baby and the length of the ileum at birth and during the first two weeks. An approximation of the surface of the ileum was calculated using the average minimum and maximum ileum lengths and a mean diameter of 1.39 mm for 3.5 kg neonate aged 37-41 weeks at birth [13]. The mean of daily intake was 117 mL/kg per day at birth. For the second week postpartum, breast milk intake (mL/day) was calculated according to Rios-Leyvraz M and Yao Q. [14]:

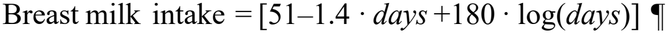

The total volume was converted into HMP weight (mg/cm^2^ ileum per day) using the amount of HMP/ml deposited onto the we **l**s (Table 2). Only a few samples of M3 from D2 to D5 follow ing birth were tested at a concentration close to the expected ingested HMP per day. The samples from the other donations were subjected to we **l**s at higher concentrations than the calculated limits to detect passage of BLps. It is remarkable that at concentrations mimicking the physiological status, passage of BLps through the ce **l** layer occurred at a very early stage, then decreased over time for similar amounts. Moreover, even assayed at higher concentrations than the calculated limits, most of the assays failed to show BLps crossing. BLps from M58 donation at D13 postpartum exhibited a positive crossing for more than 5 times the upper calculated limits. It was even worse at D15 postpartum (6 pm) for the crossing was observed for 10 times the concentration of the upper calculated limit. It suggested that the BLps transfer was unlikely to occur in both cases taking into accout the breast milk intake during one breastfeeding session [15]

## Discussion

To the best of our knowledge, this is the first study investigating the occurrence of free bifidobacterial lipoproteins in breastmilk. Our first attempt using classical biochemical approaches (i.e. gel chromatography) failed to isolate BLps. Using the CVB4 affinity column, the present study demonstrated that free BLps were present in a **l** donations from 8 mothers (39 samples). Evidence that human milk contained free Blps known to neutralize ce **l** infection with CVB provided a more comprehensive vision of the anti-viral molecules supplied to the neonate. Analysis of anti-CVB4 IgA in donations from M3, M9, M22 and M58 showed a complementation in several samples with BLps when IgA dropped.

It is unlikely that the source of BLps in the donations was the bifidobacteria displayed in the samples. During *in vitro* fermentation, the maximum release of BLps is 100µg for 10^5^ cfu *B.longum*. In the present study, as shown in Fig. 4, more than 100µg BLps (mostly *B.longum* Lps) were quantified in the M3 sample collected at 7 pm on D3 but the enumeration of *B.longum* was below the threshold of 10 copies. In contrast, on D4, at 10:30 am, about 2 log *B.longum* 314 DNA copies were enumerated, whereas BLps did not comprise *B.longum* protein sequence.

One more reason to envisage another HM BLps source than a release by milk bifidobacteria is the high variability in *Bifidobacterium* species detection. A reduced abundance in the *Bifidobacterium* genus was reported in HM from 1 to 24 weeks [16]. In our work, we observed during the first two weeks postpartum a compensation between *B.breve* and *B.longum*, resulting in an almost stable bifidobacterial content of around 10^3^-10^4^ copies/ml (i.e., about 10^5^-10^6^ copies/kg body weight for a mean intake of 100ml HM/kg). Nonetheless, counts per species were highly varied as demonstrated in the multiple donations. Besides, *B.longum* BLps were detected in samples lacking the corresponding bifidobacterial species. And the reverse was true. Thus, it was unlikely that HM bifidobacteria released their ce **l** wa **l** lipoproteins in the mammary gland. Overa **l**, our results suggested an extramammary source of BLps unrelated to the milk bifidobacteria.

One might assume that the main source of milk BLps is the intestine, since bifidobacteria highly colonize the human gut, primarily the ascending colon, where they can release their lipoproteins. However, the source of milk bifidobacteria themselves is sti **l** a matter of debate.

Following the establishment of bifidobacteria in the infant gut, both the entero-mammary route and the cross-contamination with bifidobacteria during the baby’s suckling can contribute to the milk contamination with bifidobacteria [17-20]. In our study, we noted that in the very first hours following birth, the first milk secretions can harbour bifidobacterial DNA. Of note, bifidobacte ria are established at a high level in the term breastfed infant gut within the 4^th^ day postpartum. Thus, bifidobacterial detection in HM donations at a very early stage following delivery, before gut establishment, corroborated the possible translocation of bifidobacteria from the mother’s intestine [21]. The translocation of bifidobacteria implied that they were close to the intestinal ce **l**s. The uptake of their ce **l** wa **l** Lps once released in the intestinal environment could be facilitated by the close contact between bacteria and ce **l**s. To substantiate this assumption, BLps isolation from stools could help confirm the relevance of the mother’s intestine as the source of BLps, although the localisation of BLps releasing bifidobacteria is primarily the distal ileum and the ascending colon.

Once free BLps were detected in HM, we wondered about the fate of DHM BLps in the newborn’s intestine, especia **l**y considering that the target of CV B is the pancreas. The key issue was to determine the extent to which the DHM BLps could reach the internal organs to delineate the potential of HM BLps to prevent the systemic spread of coxsackievirus. We have already investigated the ability of BLps isolated from *B. longum* fermentation to reach the internal organs *in vivo* [22]. TRITC labe **l**ed BLps were not recovered in Peyer’s patches and spleen, contrary to intestinal bacteria. Instead, the labelled BLps were detected in bone marrow. It suggested another route than the bacterial uptake by M ce **l**s. Flux across the intestinal epithelial barrier occurs via transepithelial transport, comprising transcellular and parace **l**ular pathways. A possible passage via the epithelial barrier was addressed using the cultured human epithelial Caco-2 ce **l**s grown onto a 0.4 μm pore size of the insert’s membrane that prevented the passage of any bacteria, comprising bifidobacteria. Thus, a positive crossing of DHM BLps supported a mechanis m unrelated to bacterial pathways.

Flux across the epithelial barrier occurs via transepithelial transport, comprising transce **l**ular and parace **l**ular pathways. Transcellular transport involves movement of molecules through ce **l**s and is mediated by apical and basolateral transmembrane transporters with exquisite substrate specificity. Parace **l**ular transport is less selective and can involve movement of molecules across the epithelial barrier via the pore pathway or the leak pathway. The tight junction leak pathway a **l**ows molecules with diameters up to ∼12.5 nm to traverse the epithelial barrier whereas the maximum diameter of solutes that can pass through channels generated by pore-forming claudins is 0.6 nm [23]. It was dubious that DHM BLps were crossing the ce **l** layer through the pore or leak pathways, considering the size of either the aggregated or the monomeric forms [8, 9]. Transcellular transport was more likely the pathway for BLps to cross the ce **l** layer. MS analysis of the 80 kDa bands detected by SDS-PAGE in the lower compartment after 4h incubation of the Caco2 ce **l** layer with M17 DHM samples (D5 and D7) distinguished intact lactoferrin contaminating BLps. Human lactoferrin was previously shown to be taken up into the 21 d-cultured Caco2 ce **l**s and released intact in the culture medium thanks to the lactoferrin receptor, the intelectin-1 (Itln1), present in the Caco2 ce **l**. The intelectin-1, expressed in Caco-2 ce **l** line, is a disulfide-bonded, N-glycosylated, trimeric, secreted protein belonging to a family of calcium-dependent lectins which recognize a set of microbial glycans [24-26]. Our previous studies identified carbohydrates in the fractions comprising the BLps [8, 9]. Bionyc search through the BLps peptides released following tryptic digestion substantiated the O-linkage of the oligosaccharides to the protein sequence. It implied that the O-glycosylated BLps could act as a ligand of the ce **l** lectins. Galectin-1 was already shown to bind BLps from *B.longum* and *B.breve*. Since Gal-1 is not expressed in Caco-2 ce **l** line, the intelectin-1 expressed in the ce **l** line and recognizing galactose could conceivably be the receptor candidate [27, 28].

Lastly, even if the BLps crossing can occur, the volume consumed by the neonate could limit the passage. Evidence was provided that BLps passage was compatible with colostrum intake within the first 5 days following birth. Later, the amount of milk powder to apply to the ce **l** model for a positive crossing was not compatible with a standard intake. However, the 21 d-cultured Caco-2 ce **l**s do not reproduce the intestinal permeability of the immature gut which likely facilitate the process [29-33]. Besides, other HM components, i.e. the discarded oligosaccharides could also impact the rate of transfer. HM neutral oligosaccharides were shown to be transported across the intestinal epithelium by receptor-mediated transcytosis and by parace **l**ular pathways whereas acidic oligosaccharides translocated from the apical compartment to the basal one via the parace **l**ular flux [34]. To surmount the limitations, further investigation is necessary to clarify the transfer capacities of HM BLPs *in vivo*, which would indicate not just localized but also systemic antiviral protection for the infant.

## Conclusion

In conclusion, using CV B4 as a sensor uncovers a broader anti-viral repertoire in donor human milk encompassing bifidobacterial lipoproteins alongside IgA antibodies. It underscored significant fluctuations in the BLps amounts and qualities each day and within a day, setting a foundation for new strategies in infant food supplement formulations. Additional research is, however, needed to decrypt the HM BLps diversity, to identify the nutritional factors 397 influencing HM BLps amounts and to assess the distribution of intestinal BLps in mothers.

Further exploration into the BLps transport is also necessary to determine the degree of any possible systemic protection against coxsackievirus.

## Acknowledgments

The authors wish to thank Dr. Véronique Pierrat, the biobank staff and the participants for their contributions. The authors would also like to thank Frédéric Huguet and Sokunthéany Verron-Ly from Bifinove SAS for their technical assistance.

## Author contributions

**Siwar Ayari, Fabrice Bray, Famara Sané**, and **Stéphanie Devassine:** Substantial contribution to acquisition of data, analysis and interpretation of data, drafting the article.

**Marie-Bénédicte Romond:** Substantial contribution to conception and design, drafting the article and Final approval of the version to be published.

**Frank Piva, Philippe Gervois:** Substantial contribution to drafting the article, revising it critica lly and Final approval of the version to be published.

A **l** authors approved the final manuscript as submitted and agree to be accountable for a **l** aspects of the work.

## Consent statement

Information about the study was given to the mothers wi **l**ing to donate their milk to the biobank. The donors motivated to participate to the study were asked for their written consent before being enrolled in the study.

## Data Availability

Data can be shared on demand with the exception of personal data

## References

1. J, Carroll, Chen J, Mittal R, Lemos JRN, Mittal M, Juneja S, Assayed A and Hirani K, Decoding the Significance of Alpha Cell Function in the Pathophysiology of Type 1 Diabetes. Ce ls, 2024. 13(22): p. 1914.

2. A, Turin, Dovč K, Klemenčič S, Bratina N, Battelino T, Lipovšek JK, Uršič K, Shmueli-406 Goetz Y and Drobnič-Radobuljac M, Carer’s Attachment Anxiety, Stressful Life-Events and the Risk of Childhood-Onset Type 1 Diabetes. Front Psychiatry, 2021. 12: p. 657892.

3. A, Carré, Vecchio F, Flodström-Tullberg M, You S and Ma lone R, Coxsackievirus and Type 1 Diabetes: Diabetogenic Mechanisms and Implications for Prevention. Endocr 411 Rev, 2023. 44(4): p. 737–751.

4. MP, Nekoua, Mercier A, Vergez I, Morvan C, Mbani CJ, Sane F, Lobert D, Engelmann I, Romond MB, Alidjinou EK and Hober D, Infection à coxsackievirus B et pathogenèse du diabète de type 1 [Coxsackievirus B infection and pathogenesis of type 1 diabetes]. Virology, 2022. 26(6): p. 415–430.

5. F, Sane, Alidjinou EK, Kacet N, Moukassa D, Charlet C, Ebatetou-Ataboho E, Ngoulou W, Badia-Boungou F, Romond MB and Hober D, Human milk can neutralize 418 Coxsackievirus B4 in vitro. . J Med Virol, 2013. 85(5): p. 880–887.

6. KA El Kfoury, Romond MB, Scuotto A, Alidjinou EK, Dabboussi F, Hamze M, Engelmann I, Sane F and Hober D, Bifidobacteria-derived lipoproteins inhibit infection with coxsackievirus B4 in vitro. Int J Antimicrob Agents, 2017. 50(2): p. 177–185.

7. R. Martín, Jiménez E, Heilig H, Fernández L, Marín ML, Zoetendal EG and Rodríguez JM, Isolation of bifidobacteria from breast milk and assessment of the bifidobacterial population by PCR-denaturing gradient gel electrophoresis and quantitative real-time PCR. . Appl Environ Microbiol., 2009. 75(4): p. 965–969.

8. A, Scuotto, Djorie S, Colavizza M, Romond PC and Romond MB, Bifidobacterium breve C50 secretes lipoprotein with CHAP domain recognized in aggregated form by TLR2. Biochimie, 2014. 107: p. 367–375.

9. A, Scuotto, Romond PC, Djorie S, Alric M and Romond MB, In silico mining and characterization of bifidobacterial lipoprotein with CHAP domain secreted in an aggregated form. . Int J Biol Macromol, 2016. 82: p. 653–662.

10. F, Sane, Scuotto A, Pierrat V, Kacet N, Hober D and Romond MB, Diabetes progression and alterations in gut bacterial translocation: prevention by diet 434 supplementation with human milk in NOD mice. J Nutr Biochem, 2018. 62: p. 108–122.

11. V, Cleusix, Lacroix C, Dasen G, Leo M and Le Blay G, Comparative study of a new quantitative real-time PCR targeting the xylulose-5-phosphate/fructose-6-phosphate phosphoketolase bifidobacterial gene (xfp) in faecal samples with two fluorescence in situ hybridization methods. J Appl Microbiol, 2010. 108(1): p. 181–193.

12. J, Gao, Hugger ED, Beck-Westermeyer MS and Borchardt RT, Estimating intestinal mucosal permeation of compounds using Caco-2 cell monolayers. Curr Protoc Pharmacol, 2001. 7(2).

13. C, Bardwel, El Deme lawy D, Oltean I, Murphy M, Agarwal A, Hamid JS, Reddy D, Barrowman N, de Nanassy J and Nasr A, Establishing normal ranges for fetal and neonatal small and large intestinal lengths: results from a prospective postmortem 445 study. World J Pediatr Surg, 2022. 5(3).

14. M, Rios-Leyvraz and Yao Q, The Volume of Breast Milk Intake in Infants and Young 447 Children: A Systematic Review and Meta-Analysis. Breastfeed Med, 2023. 18(3): p. 188–197.

15. L, Gu, Dong L and Chen H, Relationships of breast milk intake during one breastfeeding 450 session with sucking time and other determinants: a cross-sectional study. Int Breastfeed J, 2025. 20(1): p. 70.

16. KE, Lyons, Shea CO, Grimaud G, Ryan CA, Dempsey E, Ke ly AL, Ross RP and Stanton C, The human milk microbiome aligns with lactation stage and not birth mode. Sci Rep, 2022. 12(1).

17. T, Jost, Lacroix C, Braegger CP, Rochat F and Chassard C, Vertical mother-neonate transfer of maternal gut bacteria via breastfeeding. Environ Microbiol, 2014. 16(9): p. 2891–2904.

18. K, Kordy, Gaufin T, Mwangi M, Li F, Cerini C, Lee DJ, Adisetiyo H, Woodward C, Pannaraj PS, Tobin NH and Aldrovandi GM, Contributions to human breast milk microbiome and enteromammary transfer of Bifidobacterium breve. PLoS One, 2020. 15(1).

19. Y, Du, Qiu Q, Cheng J, Huang Z, Xie R, Wang L, Wang X, Han Z and Jin G, Comparative study on the microbiota of colostrum and nipple skin from lactating 464 mothers separated from their newborn at birth in China. Front Microbiol, 2022. 13.

20. C, Qi, Zhou J, Tu H, Tu R, Chang H, Chen J, Li D, Sun J and Yu R, Lactation-dependent vertical transmission of natural probiotics from the mother to the infant gut through 467 breast milk. Food Funct, 2022. 13(1): p. 304–315.

21. L, Ruiz, Bacigalupe R, García-Carral C, Boix-Amoros A, Argüello H, Silva CB, de Los Angeles Checa M, Mira A and Rodríguez JM, Microbiota of human precolostrum and its potential role as a source of bacteria to the infant mouth. Sci Rep, 2019. 9(1).

22. F, Piva, Gervois P, Karrout Y, Sané F and Romond MB, Gut-Joint Axis: Impact of Bifidobacterial Cell Wall Lipoproteins on Arthritis Development. Nutrients, 2023. 15(23): p. 4861.

23. A, Horowitz, Chanez-Paredes SD, Haest X and Turner JR, Paracellular permeability and tight junction regulation in gut health and disease. Nat Rev Gastroenterol Hepatol, 2023. 20(7): p. 417–432.

24. T, Matsuzaki, Nakamura M, Nogita T and Sato A, Cellular Uptake and Release of Intact Lactoferrin and Its Derivatives in an Intestinal Enterocyte Model of Caco-2 Cells. Biol Pharm Bu l, 2019. 42(6): p. 989–995.

25. Y, Akiyama, Oshima K, Shin K, Wakabayashi H, Abe F, Nadano D and Matsuda T, Intracellular retention and subsequent release of bovine milk lactoferrin taken up by human enterocyte-like cell lines, Caco-2, C2BBe1 and HT-29. Biosci Biotechnol 483 Biochem, 2013. 77(5): p. 1023–1029.

26. DA, Wesener, Wangkanont K, McBride R, Song X, Kraft MB, Hodges HL, Zarling LC, Splain RA, Smith DF, Cummings RD, Paulson JC, Forest KT and Kiessling LL, Recognition of microbial glycans by human intelectin-1. Nat Struct Mol Biol, 2015. 22(8): p. 603–610.

27. BT, Maybruck, Pfannenstiel LW, Diaz-Montero M and Gastman BR, Tumor-derived exosomes induce CD8+ T cell suppressors. J Immunother Cancer, 2017. 5(1): p. 65. S, Sharma and Ramya TNC, Saccharide binding by intelectins. Int J Biol Macromol, 2018. 108:p. 1010–1016.

28. JO, Gebbers and Laissue JA, Bacterial translocation in the normal human appendix parallels the development of the local immune system. Ann N Y Acad Sci, 2004. 1029: p. 337–343.

29. L, Geurts, Neyrinck AM, Delzenne NM, Knauf C and Cani PD, Gut microbiota controls adipose tissue expansion, gut barrier and glucose metabolism: novel insights into molecular targets and interventions using prebiotics. Benef Microbes, 2014. 5(1): p. 3–17.

30. JY, Yoo, Groer M, Dutra SVO, Sarkar A and McSkimming DI, Gut Microbiota and 500 Immune System Interactions. Microorganisms, 2020. 8(10): p. 1587.

31. JCW de Jong, Ijssennagger N and van Mil SWC, Breast milk nutrients driving intestinal epithelial layer maturation via Wnt and Notch signaling: Implications for necrotizing enterocolitis. Biochim Biophys Acta Mol Basis Dis, 2021. 1867(11): p. 166229.

32. KM, Robertson, Cermack K, Woodruff T, Carter SR, Jala VR and Buonpane C, Review: microbial metabolites -a key to address gut inflammation and barrier dysfunction in the premature infant. Gut Microbes., 2025. 17(1).

33. MJ, Gnoth, Rudloff S, Kunz C and Kinne RK, Investigations of the in vitro transport of human milk oligosaccharides by a Caco-2 monolayer using a novel high-performance liquid chromatography-mass spectrometry technique. J Biol Chem, 2001. 276(37): p. 34363–34370.

