## Supplemental Table S1 for "Detection of bifidobacterial lipoproteins with anti-viral potential in donor human milk"

Table S1: Detection of *B. breve* and *B. longum* during the 2^nd^ week postpartum.

| Day (postpartum) | Mother | *B.longum* (log cfu/ml) | *B.breve* (log cfu/ml) |
| --- | --- | --- | --- |
| 7 | 29 | ND | ND |
| 8 | 7 | ND | 2.3 |
| 9 | 30 | ND | 3.5 |
| 10 | 53 | ND | ND |
|  | 57 | ND | 3.5 |
| 11 | 42 | ND | ND |
| 13 | 54 | ND | 3.3 |
| 14 | 14 | ND | 2.8 |
| 15 | 23 | ND | 2.9 |

**ND :** Not detected
