## Supplemental Table S2 for "Detection of bifidobacterial lipoproteins with anti-viral potential in donor human milk"

Table S2 : Additional titration of BLps and anti-CV B4 IgA

| Donor | Donation (days post-partum) | BLps (µg/mg HMP) | *B. longum* (log10 cfu/ml) | *B.breve* (log10cfu/ml) | anti-CV B4 IgA (AU/mg HMP)^a^ |
| --- | --- | --- | --- | --- | --- |
| 24 | D4 03:40 pm | 0.6 | ND | ND | 6.85 |
| 22 | D6 06:40 pm | 2.5 | 1.24 | ND | ND |
|  | D6 11:00pm | 5.0 | 1.21 | ND | 4.7 |
|  | D7 10:40 am | 2.5 | ND | ND | 10.45 |
| 29 | D8 00:20 am | 2.5 | 2.36 | 1.76 | 3.48 |
| 9 | D10 05:00 pm | 0.6 | 2.44 | ND | ND |
|  | D10 07:20 pm | 2.5 | ND | ND | ND |
|  | D10 11:30 pm | 2.5 | ND | ND | 4.4 |

^a^: HMP exhibiting the highest neutralizing titer (i.e 8AU/mg HMP) against CV B4 infection was used for the standard curve. Results are expressed as arbitrarily unit (AU)/mg HMP.
