## Supplementary figures and images for "Detection of bifidobacterial lipoproteins with anti-viral potential in donor human milk"

### Supplemental Table S3

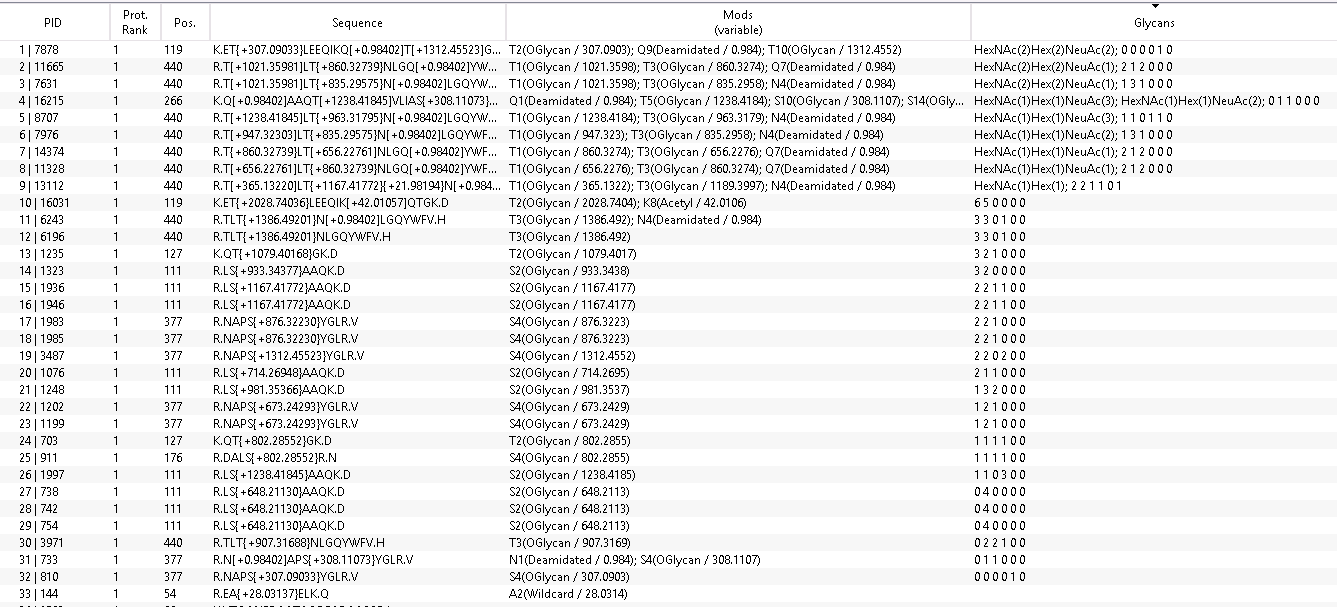
Table S3: Byonic analysis of *B.longum* CBi0703 peptides released by trypsin digestion
